# Orthogonal cell division organizes surface virulence factors to drive staphylococcal abscess community formation

**DOI:** 10.64898/2026.03.12.711374

**Authors:** Félix Ramos-León, Valerie Altouma, Peri Goldberg, Amany M. Ibrahim, Domenico D’Atri, Federico Machinandiarena, Vicenzo Verdi, Dominique M. Missiakas, Kimberly M. Davis, Kumaran S. Ramamurthi

## Abstract

During infection, *Staphylococcus aureus* forms dense multicellular structures, called staphylococcal abscess communities (SACs), that are encased in a capsule made of host fibrin to evade host immune defenses. *S. aureus* cells divide characteristically along successive orthogonal planes, but the contribution of this division geometry to infection is unclear. Here, we show that disrupting orthogonal cell division by deleting the cell division septum placement factor PcdA impairs SAC formation *in vivo* and in a three-dimensional *in vitro* model. Loss of PcdA leads to uneven surface distribution of adhesins containing the YSIRK signal sequence that directs their insertion into the division septum, thereby resulting in uneven interaction with fibrin fibers. Consequently, bacterial communities fail to establish a robust fibrin pseudocapsule and remain accessible to immune cells. We propose that orthogonal cell division coordinates cell cycle progression with extracellular matrix engagement, SAC architecture, and persistence within host tissues.

**HIGHLIGHTS:**

- Orthogonal cell division promotes staphylococcal abscess community formation
- Loss of PcdA disrupts fibrin pseudocapsule assembly in 3D models
- Division geometry ensures uniform surface deployment of adhesins
- Cell division plane selection links bacterial cell cycle control to virulence

## INTRODUCTION

*Staphylococcus aureus* is a Gram-positive commensal of the human skin and nasal cavity that is also an opportunistic pathogen and is a leading cause of nosocomial invasive infections, including soft tissue infections, osteomyelitis, and endocarditis ^1^. *S. aureus* cells are spherical and divide by binary fission in characteristic consecutive orthogonal planes ^2–4^. Unlike rod-shaped bacteria, which rely on pre-existing polarity cues, *S. aureus* divides without obvious intrinsic spatial landmarks. Placement of the division plane is controlled by two independent pathways. One pathway depends on the NTPase PcdA (conserved among staphylococci and closely related species), which recruits the central cell division protein FtsZ to the division site via direct interaction ^5^. PcdA identifies the correct division plane by interacting with the structural protein DivIVA, which preferentially embeds into membranes of increased micron-scale negative curvature ^5,6^. Although *S. aureus* is typically spherical (and therefore would not display regions of increased membrane curvature), the PBP3-RodA cell wall synthesis machinery drives a subtle elongation of *S. aureus* cells during growth to generate a slightly ovoid morphology and therefore a transient axial asymmetry ^7^, which DivIVA exploits to mark the next division plane ^5^. The second pathway, independent of PcdA, invokes the phenomenon of nucleoid occlusion, whereby the Noc protein, in addition to controlling DNA replication initiation, is proposed to inhibit FtsZ polymerization over the chromosome and abrogates orthogonal cell division ^5,8,9^.

In the absence of PcdA, cells display increased sensitivity to cell wall-targeting antibiotics ^5,10^ and reduced virulence in a murine renal abscess model of infection ^5^. Moreover, *pcdA* was also identified in clinical *S. aureus* isolates from severe infections as accumulating “adaptive” mutations that may permit increased persistence in a host ^11^. Despite these phenotypes, functional consequences of cell division plane selection during infection remain unclear.

During invasive intravenous infection in a mouse model, *S. aureus* forms kidney abscesses that contain a dense central bacterial core, known as a staphylococcal abscess community (SAC), surrounded by layers of necrotic and viable immune cells (predominantly neutrophils with an exterior macrophage ring) ^12,13^. This structured mode of growth restricts bacterial dissemination while promoting persistence within host tissues. Genetic analyses have identified multiple determinants required for SAC formation, many of which are established virulence factors ^12–15^. A critical step in this process is the conversion of host fibrinogen into fibrin through the activity of coagulases secreted by *S. aureus* ^13,16^. *S. aureus* organizes these fibrin fibers into a defined outer layer, termed the pseudocapsule, that protects the bacterial community from the surrounding immune cells ^13,16^. Thus, bacterial cell surface interactions with fibrin and fibrinogen are central to abscess development. Interaction with fibrin and fibrinogen is mediated by cell wall-anchored adhesins (also known as microbial surface components recognizing adhesive matrix molecules (MSCRAMMs)), including clumping factor A (ClfA), fibronectin-binding proteins A and B (FnbpA/b), and SrdE ^17,18^. These proteins share a conserved architecture, consisting of an N-terminal signal sequence, extracellular domains responsible for ligand binding, and a C-terminal LPXTG sorting signal that is recognized by sortase A for covalent attachment to the peptidoglycan ^17,19^. Notably, many of these adhesins contain a conserved YSIRK-GXXS motif within their signal sequence, which targets their secretion to the division septum ^20,21^. Recent studies have demonstrated that secretion of YSIRK-containing proteins is tightly coupled to septal cell wall synthesis, thereby linking surface protein localization directly to the bacterial cell cycle ^21–24^.

In this study, we investigate how regulation of division plane selection contributes to abscess community formation in *S. aureus*. Using a mouse infection model together with host-relevant three-dimensional models of SAC growth, we show that PcdA-dependent orthogonal cell division is required for proper SAC architecture and fibrin pseudocapsule assembly. We further demonstrate that disruption of division plane selection alters interaction with fibrin by affecting the surface distribution of adhesins that harbor a YSRIK signal sequence, thereby linking orthogonal cell division and cellular organization to abscess development and pathogenesis.

## RESULTS

### Deletion of pcdA impairs establishment of staphylococcal abscess communities (SACs) in vivo

We previously showed, using a murine renal abscess model of infection, that mice infected with a Δ*pcdA* strain developed abscesses with reduced detectable bacterial growth compared to WT infections ^5^. Additionally, although infection by both WT and Δ*pcdA* strains produced a similar number of abscesses by day 5, at day 15, the number of abscesses in kidneys from the Δ*pcdA* infection did not increase, whereas those from the WT infection continued to increase. Together, this suggested that deletion of *pcdA* may affect progression through the different stages of SAC development. To test this, we infected mice intravenously with either WT or Δ*pcdA* strain constitutively expressing the fluorescent protein mScarlet3 under control of the ribosomal *rplM* promoter. Kidneys were then harvested 3 days post-infection, fixed, and analyzed by immunofluorescence microscopy. Bacteria were visualized by mScarlet3 fluorescence, and neutrophil infiltration was detected by Hoechst staining of host cell nuclei. Distinct regions of infection within kidneys were analyzed, and bacterial growth was classified according to established criteria for the four different stages of SAC development ^14^. Stage I of SAC development is characterized by phagocytosed bacteria observed as single cells or small clusters closely associated with neutrophil nuclei (Fig. 1A-A’’). Stage I regions represented only 14% of the total areas analyzed in kidneys infected with the WT strain (29 total areas), but 34% of the areas in kidneys infected with the Δ*pcdA* strain (35 total areas). Stage II is characterized by the presence of extracellular bacterial clusters (Fig. 1B-B’’) and accounted for 24% and 23% of the areas in WT and Δ*pcdA* infections, respectively. Stage III is defined by the elaboration of mature SACs, which appear as compact bacterial masses that are spatially separated from surrounding immune cells (Fig. 1C-C’’). At this stage, mScarlet3 fluorescence was unexpectedly reduced, and bacteria were therefore identified primarily based on nucleic acid staining. Stage III regions represented 35% of the area in WT-infected kidneys, but only 17% in Δ*pcdA*-infected kidneys. Finally, Stage IV regions are characterized by SAC disruption and subsequent dispersion of bacteria (Fig. 1D-D’’), which accounted for 27% of the area in the WT infections and 26% in Δ*pcdA* infections. The increase of the fraction of Δ*pcdA* infections represented by Stage I compared to WT and the concomitant decrease in representation of Stage III suggested that Δ*pcdA* cells may be retarded in their progression through the stages of SAC development. Consistent with this notion, we observed that Stage I lesions formed by the Δ*pcdA* strain showed a reduced bacterial area compared to WT (Figure 1E), whereas no significant differences were detected between WT and the Δ*pcdA* in Stages II-IV (Fig. S1A). Similarly, the total lesion area, defined as the best-fit rectangular area encompassing the bacterial cells revealed a similar decrease in size at Stage I from lesions formed by the Δ*pcdA* strain (Fig. 1F, Fig. S1B). Finally, Stage I lesions formed by infection with the Δ*pcdA* strain resulted in reduced bacterial density (reported as the fraction of the total rectangular area occupied by bacteria) (Fig. 1G, Fig. S1C). Together, the data indicate that deletion of *pcdA* specifically alters the properties of lesions at the early stage of infection in the murine model.

**Figure 1.**
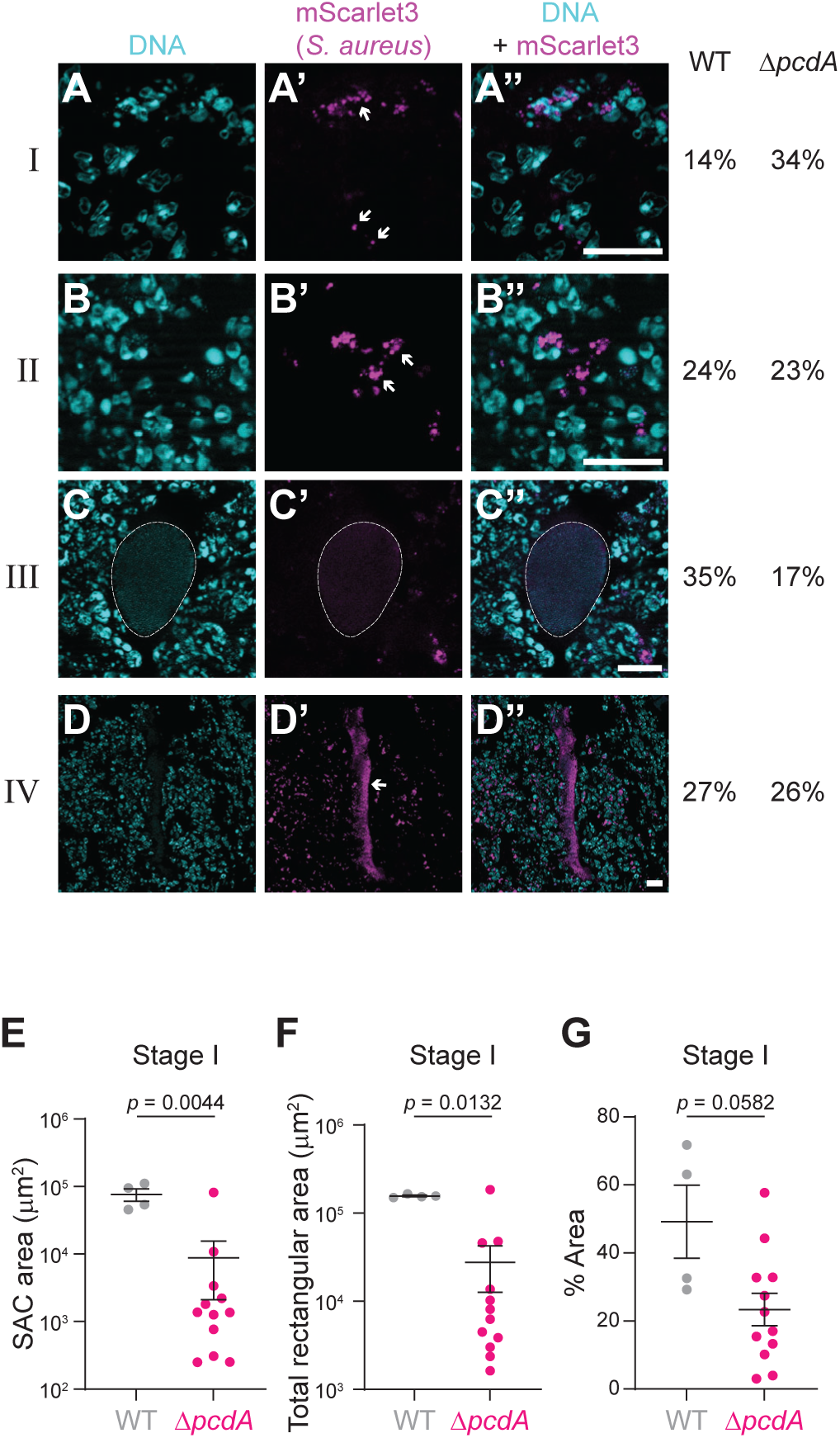
Deletion of *pcdA* impairs establishment of staphylococcal abscess communities (SACs) *in vivo*. (A-D) Representative immunofluorescence images illustrating stages I-IV of staphylococcal abscess colony formation by the WT strain. Left: DNA visualized with Hoechst stain, allowing identification of neutrophil nuclei and extracellular *S. aureus*; center: mScarlet3 fluorescence produced by *S. aureus* cells; right: overlay, Hoechst and mScarlet3. Stages of SAC formation indicated to the left. (A-A’’) Stage I, characterized by intracellular single cells or small clusters of *S. aureus* (arrows); (B-B’’) Stage II, characterized as small extracellular bacterial clusters (arrows). (C-C’’) Stage III represents well-established SACs; the SAC perimeter is indicated by a dotted line. (D-D’’) Stage IV, characterized by disrupted SACs (arrow). The percentage of each infection stage observed in kidneys infected with the WT or the Δ*pcdA* strain is shown on the right. Scale bars: 20 µm (note the different magnifications in C-D). (E-G) Quantification of the following parameters of cells in Stage I lesions in kidneys of mice infected with WT (grey) and Δ*pcdA* (pink): (E) SAC area (area occupied by bacterial cells), (F) total rectangular area (area of a best-fit rectangle around the bacterial cells), and (G) % area (a density measurement that reports the % of the best-fit rectangle that is occupied by bacterial cells). Bars indicate mean; errors: S.D. *P* values were determined using a Mann-Whitney test.

### *pcdA* deletion does not alter bacterial uptake by phagocytic cells

The increased proportion of stage I lesions led us to assess if *pcdA* deletion alters the susceptibility of individual cells to phagocytic uptake. We therefore performed bacterial uptake assays using the RAW264.7 murine macrophage cell line and *S. aureus* cells expressing mScarlet3 from the constitutive *rplM* promoter. WT and Δ*pcdA* cells were added to macrophages at a multiplicity of infection (MOI) of 5 and incubated for 2 hours to allow uptake. Following incubation, extracellular bacteria were eliminated by gentamicin treatment, and macrophage nuclei were stained with Hoechst and their periphery with fluorescently labeled wheat germ agglutinin. Using fluorescence microscopy, we observed bacteria as small intracellular clusters within macrophages (Fig. 2A-B’’’). Quantification of the percentage of macrophages containing intracellular bacteria revealed no significant difference between infections with the WT strain (27.4% ± 8.5) and the Δ*pcdA* strain (31.7% ± 4.6) (Fig. 2C). To validate these observations using an independent approach, the infection assay was also analyzed by flow cytometry (Fig. 2D, Fig. S2). Consistent with the microscopy-based quantification, similar proportions of infected cells were observed for the WT (27.5% ± 8.6) and Δ*pcdA* (28.9% ± 5.0) strains, indicating that deletion of *pcdA* does not increase uptake by phagocytic cells.

**Figure 2.**
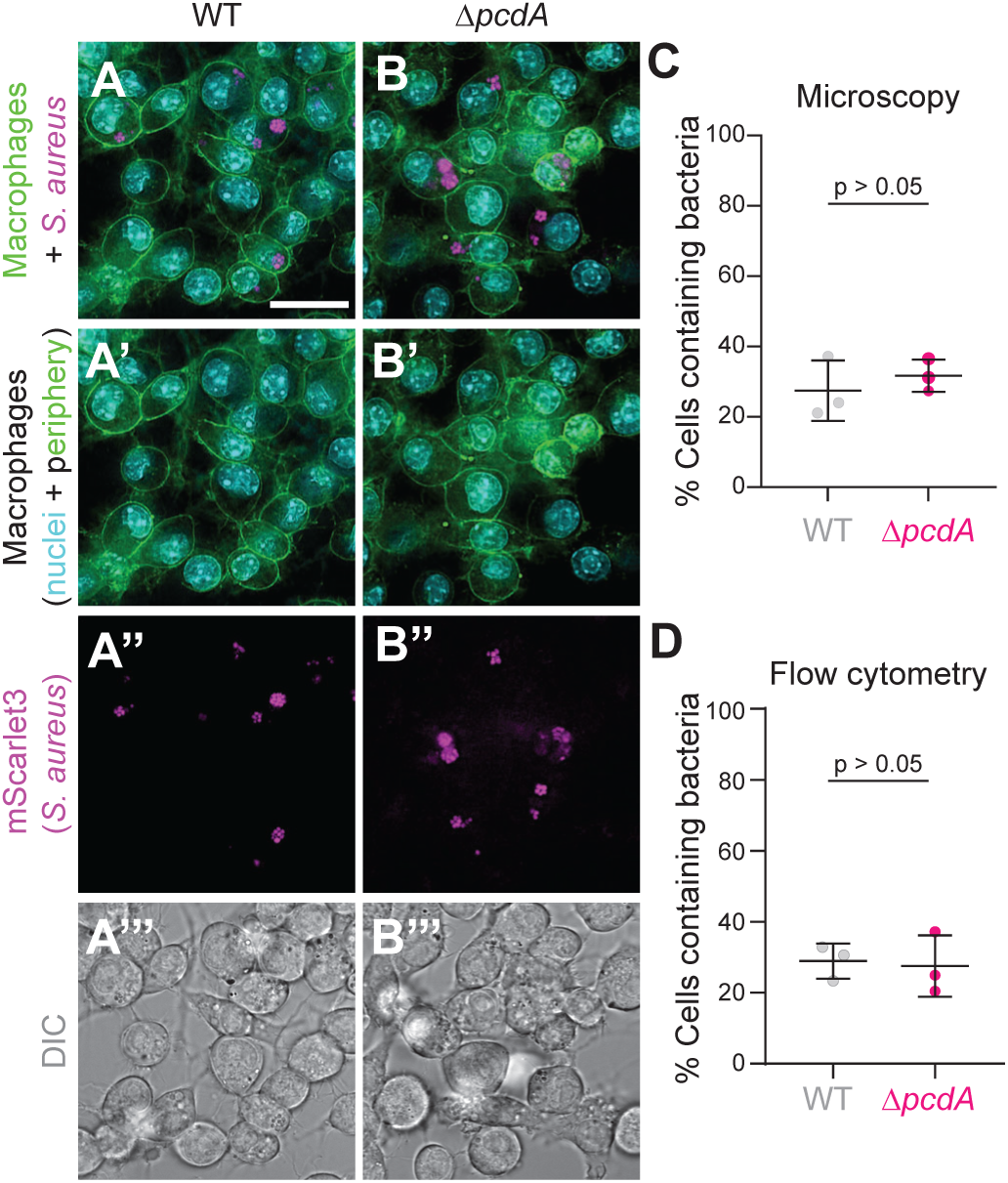
Absence of PcdA does not increase sensitivity of individual *S. aureus* cells to phagocytosis. (A-B) Fluorescence micrographs of RAW264.7 macrophages containing phagocytosed (A-A’’’) WT or (B-B’’’) Δ*pcdA S. aureus*. Macrophage cell periphery was visualized using WGA (green), and nuclei visualized using Hoechst stain (cyan); *S. aureus* cells visualized via production of mScarlet3. (A, B) Overlay, WGA, Hoechst, and mScarlet3; (A’, B’) overlay, WGA and Hoechst; (A’’, B’’) fluorescence from mScarlet3; (A’’’, B’’’) differential interference contrast (DIC). (C) Quantification of the percentage of macrophages containing intracellular bacteria based on microscopy analysis. Each independent replicate (n = 3) represents >700 cells. (D) Quantification of the percentage of macrophages containing intracellular bacteria measured by flow cytometry. Each independent replicate (n = 3) represents >40,000 cells. Bars represent mean; errors: S.D. *P* values were calculated using an unpaired *t* test. Scale bar, 20 µm.

### PcdA is required for SAC development in a 3D in vitro model

To determine whether the reduction in stage III observed *in vivo* reflects a defect in SAC formation by the Δ*pcdA* strain, we examined SAC development using a modified 3D collagen gel model supplemented with fibrinogen and prothrombin ^15,25^. WT and Δ*pcdA* strains were inoculated at low density to ensure that each SAC originated from a single bacterial cell. Under these conditions, the WT strain formed roughly spherical SACs composed of compact bacterial microcolonies with well-defined boundaries (Fig. 3A). These SACs displayed a mean area of 594.9 μm^2^ ± 322.6 (n = 178) (Fig. 3G) and an aspect ratio of 1.160 ± 0.127 (Fig. 3H). In contrast, the Δ*pcdA* strain formed SACs with pronounced morphological defects (Fig. 3B) that were smaller (reduced mean area of 95.4 μm^2^ ± 374.6 (n = 115)) and more irregularly shaped (increased aspect ratio of 1.630 ± 0.512) compared to WT SACs (Figure 3G-H). Expression of *pcdA* from a different chromosomal locus restored WT-like SAC morphology (Figure 3C), indicating that the observed defects were due to the absence of PcdA. This SAC formation defect in the Δ*pcdA* strain was not due to impaired coagulation ^16^, since cultures of the Δ*pcdA* strain retained the ability to coagulate fibrinogen, in contrast to a coagulase production-defective mutant (Δ*saeR*) that did not (Fig. S3). Deletion of *pbp3*, the cell wall transpeptidase required for cell elongation during *S. aureus* growth, results in cells that remain largely spherical during the cell cycle ^7^ and consequently fail to divide along sequential orthogonal division planes ^5^. As an independent assessment of the link between orthogonal cell division and SAC formation, we observed that the Δ*pbp3* mutant also formed defective SACs similar to the *pcdA* deletion mutant (Fig. 3D) that exhibited a mean area of 365.0 μm^2^ ± 346.3 (n = 109) and an aspect ratio of 1.458 ± 0.390 (Figure 3G-H). Interestingly, deleting *noc*, which impairs initiation of DNA replication ^8^ and whose deletion also results in a defect in orthogonal cell division, did not disrupt SAC formation, resulting in ∼WT-like SACs with mean area of 582.4 μm^2^ ± 211.4 (n = 104) and an aspect ratio of 1.167 ± 0.131 (Figure 3G-H). Finally, the Δ*saeR* mutant with reduced coagulase activity did not appreciably form SACs (Fig. 3F). The data therefore indicate that SAC development depends on cell wall-related pathways that contribute to orthogonal cell division, whereas the chromosome-dependent pathway is dispensable for correct SAC morphology. Another formal possibility is that deletion of *noc* does not result in an orthogonal cell division defect in the collagen matrix growth condition.

**Figure 3.**
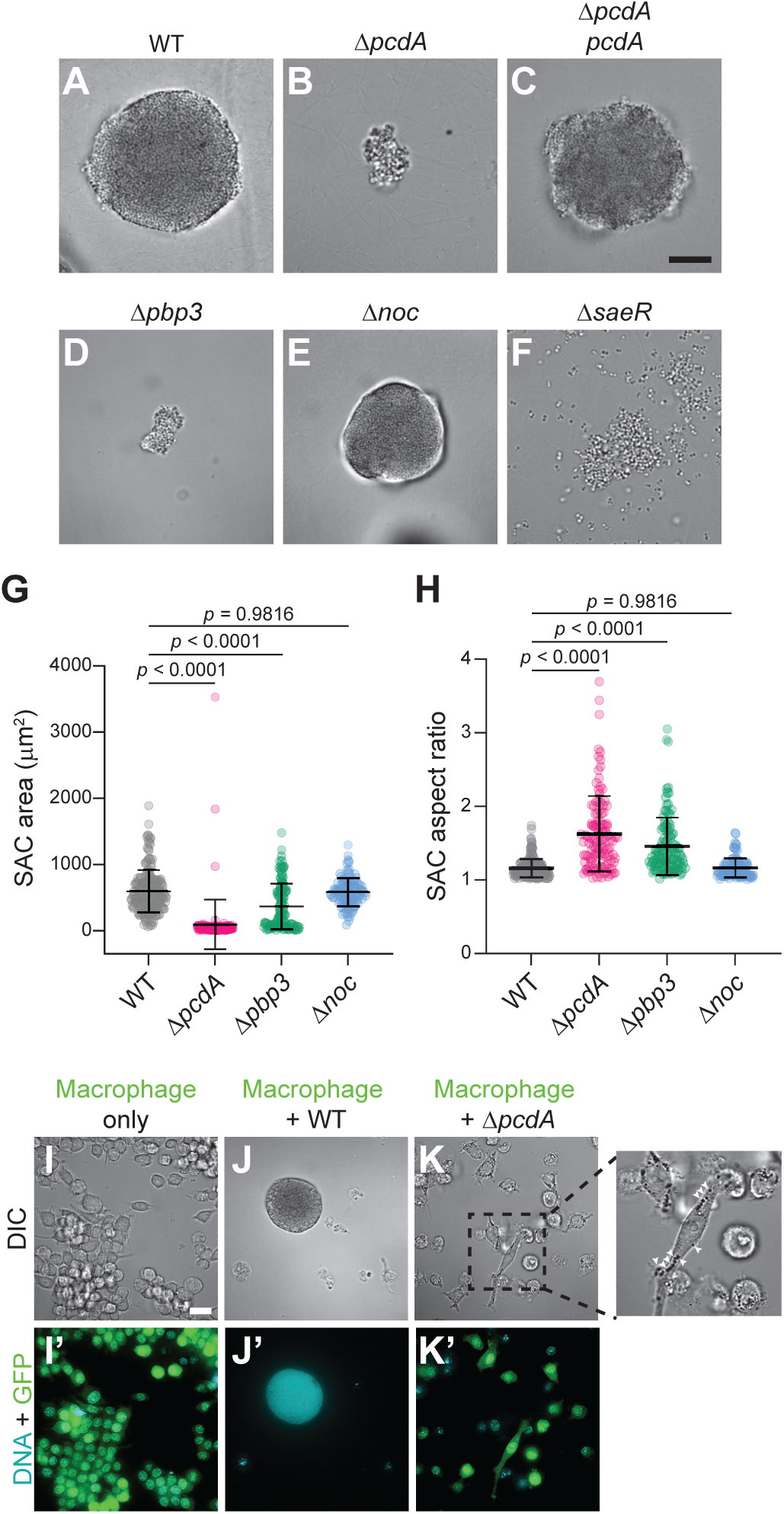
Deletion of *pcdA* disrupts SAC development in a 3D *in vitro* model. (A-F) Micrographs of *in vitro* SAC formation by (A) WT, (B) Δ*pcdA*, (C) Δ*pcdA* complemented at an ectopic genomic locus with *pcdA*, (D) Δ*pbp3*, (E) Δ*noc*, or (F) Δ*saeR* in a 3D collagen/fibrinogen/prothrombin gel. Scale bar: 20 µm. (G-H) Quantification of (D) area or (E) aspect ratio of individual SACs formed by WT (gray), Δ*pcdA* (pink), Δ*pbp3* (green), or Δ*noc* (blue). Bars indicate mean; errors: S.D. *P* values were calculated using an unpaired *t* test. (I-K’) Micrographs of 3D collagen gels containing (I, I’) only RAW264.7 murine macrophages producing GFP, or macrophages incubated with either (J, J’) WT or (K, K’) Δ*pcdA*. (I-K) DIC; (I’-K’) overlay, Hoechst stain (to visualize SACs and macrophage nuclei) and fluorescence from GFP (macrophages). Inset in (K) highlights an elongated macrophage containing internalized bacteria (white arrowheads). Scale bar: 20 µm.

We next assessed if the defective SACs formed by the Δ*pcdA* strain are more susceptible to immune clearance by incorporating cultured RAW264.7 macrophages producing EGFP (to aid in visualization) into the in vitro SAC assay. After an overnight incubation in the matrix, fluorescence microscopy revealed that, in the absence of bacteria, macrophages displayed a rounded morphology, robust EGFP fluorescence, and intact round nuclei as visualized by Hoechst staining (Fig. 3I-I’). When macrophages were co-incubated with WT *S. aureus*, we readily detected mature SACs and observed that macrophages were largely excluded from the immediate vicinity of these SACs (Fig. 3J). Those macrophages rarely observed adjacent to SACs frequently lacked detectable EGFP fluorescence and harbored poorly stained or fragmented nuclei (Fig. 3J’), consistent with a loss of membrane integrity and likely cell death. In contrast, not only did the Δ*pcdA* strain failed to form SACs, but co-incubated macrophages exhibited an elongated morphology, that frequently contained intracellular *S. aureus* cells (Fig. 3K, white arrows). Additionally, these macrophages displayed EGFP fluorescence and intact nuclear morphology, indicating that the macrophages remained viable (Fig. 3K’). Together, these observations indicate that loss of *pcdA* results in bacterial communities that are accessible to phagocytic cells in the 3D collagen model.

### PcdA is required for fibrin pseudocapsule formation and uniform fibrinogen binding

A defining feature of SACs is the fibrin pseudocapsule that physically separates enclosed bacteria from the surrounding tissue ^16,26^. To determine whether fibrin pseudocapsule formation is altered in the absence of PcdA, we incorporated fluorescently labeled fibrinogen into the 3D collagen SAC assay and analyzed fibrin organization using fluorescence microscopy. The WT strain formed SACs containing an intricate network of fibrin fibers within the bacterial community, as well as a continuous fibrin layer encapsulating the entire SAC (Figure 4A-A’). In contrast, although fluorescence was associated with Δ*pcdA* SACs, the fibrin failed to assemble into a uniform layer surrounding the community and did not form a network in the center of the SAC (Fig. 4B-B’).

**Figure 4.**
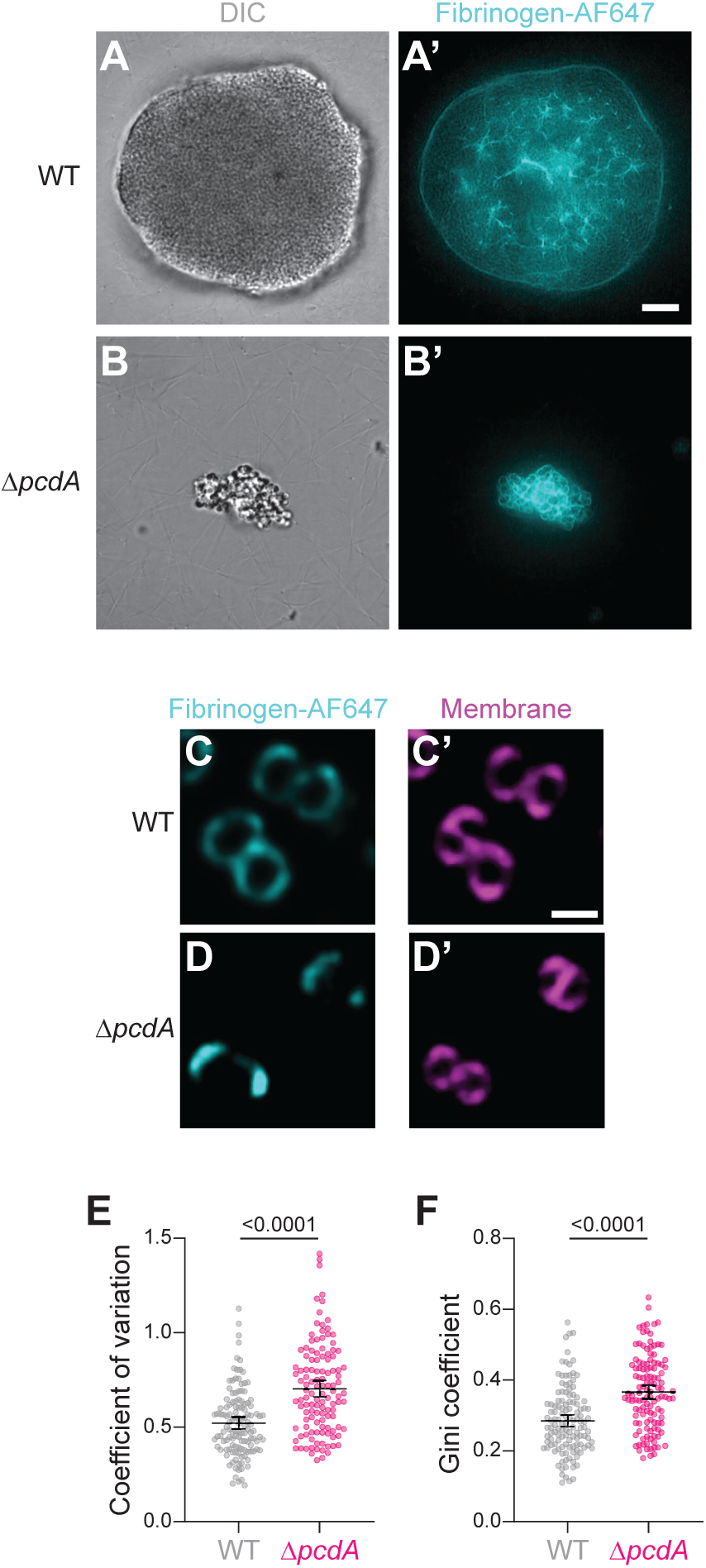
PcdA is required for fibrin pseudocapsule formation in SACs and for uniform fibrinogen binding at the single-cell level. (A-B’) Micrograph of a SAC formed by (A-A’) WT or (B-B’) Δ*pcdA* in an *in vitro* collagen gel in the presence of fluorescently labeled fibrinogen (Fibrinogen-AF647, cyan). SACs visualized by (A-B) DIC or (A’-B’) fluorescence from labeled fibrinogen. Scale bar: 20 µm. (C-D) Micrographs of individual (C-C’’) WT or (D-D’’) Δ*pcdA* cells binding fluorescently labeled fibrinogen, visualized using (C, D) membrane stain FM4-64 (C’, D’) fluorescence from labeled fibrinogen (cyan). Scale bar: 1 µm (E-F) Quantification of (E) the coefficient of variation or (F) Gini coefficient of fibrinogen fluorescence intensity around the circumference of individual WT (grey, *n* = 125) and Δ*pcdA* (pink, *n* = 120) cells. Bars represent mean ± 95% confidence interval. *P* values were calculated using a Mann-Whitney test.

To address whether this fibrin pattern reflected altered fibrinogen binding at the single-cell level, we analyzed fibrinogen binding to individual bacteria grown in a shaking culture and then labeled with fluorescently labeled fibrinogen. WT cells displayed robust and roughly homogeneously distributed fibrinogen binding around the cell periphery (Fig. 4C-C’). In contrast, Δ*pcdA* cells displayed reduced overall fibrinogen binding and the bound fibrinogen was unevenly distributed on the cell surface (Fig. 4D-D’). To quantify this relatively unequal distribution, we measured the fibrinogen fluorescence intensity along the circumference of multiple cells and calculated either the coefficient of variation (a statistical measure of data dispersion) or the Gini coefficient (a measure of inequality among the values of a frequency distribution). WT cells displayed a lower mean coefficient of variation compared to Δ*pcdA* cells (0.52 ± 0.17 (n = 125) versus 0.70 ± 0.23 (n = 120), respectively (Fig. 4E)) and mean Gini coefficient (0.28 ± 0.09 versus 0.37 ± 0.10, respectively (Fig. 4F)), indicating increased variability and unevenness in the localization of bound fibrinogen on the surface of cells in the absence of *pcdA*. We hypothesize that deletion of *pcdA* reduces SAC formation by disrupting fibrin pseudocapsule assembly, due to the altered binding and spatial distribution of fibrinogen at the surface of individual cells.

### Deletion of pcdA impairs the surface distribution of YSIRK signal sequence-containing proteins

The altered fibrinogen-binding pattern observed around Δ*pcdA* cells suggested potential defects in the surface distribution of adhesins involved in interactions with the extracellular matrix ^17^. These surface proteins are typically covalently anchored to the cell wall ^19,27^, and many of these adhesins contain a specialized signal sequence containing a YSIRK-GXXS motif that specifically targets their secretion to the division septum, although the purpose of this directed secretion and exact mechanism by which it occurs remains unclear ^20,22,28–30^. To address if deletion of *pcdA* affects the surface localization of YSIRK-containing proteins, we designed a cell surface sGFP fluorescent reporter harboring an N-terminal YSIRK signal sequence to direct septal secretion and a C-terminal cell wall sorting signal for sortase-mediated anchoring to the peptidoglycan ^31^ (Fig. 5A), expressed under control of an inducible promoter from a single chromosomal locus. 90 min after induction, we observed GFP fluorescence in WT cells, either at the division septum in cells undergoing cell division or distributed along the cell periphery in non-dividing cells (Fig. 5B-C’). In the absence of *pcdA*, in septating cells, we again observed the enrichment of fluorescence at the division septum, indicating that initial targeting of YSIRK-containing proteins to the septal region is not dependent on PcdA (Fig. 5D-E’). However, in non-dividing cells, similar to the uneven distribution of fibrinogen (Fig. 4D), we observed the relatively uneven distribution of newly synthesized surface displayed sGFP (Fig. 5D-E’). To visualize this distribution, we created a heat map of localization events in representative non-dividing cells and observed the relatively even, high fluorescence of cell surface sGFP in WT, but a patchy pattern of fluorescence in the absence of *pcdA* (Fig. 5F). This uneven distribution in the absence of *pcdA* was reflected in an increased coefficient of variation for the mutant (0.17 ± 0.04 (n = 115) versus 0.15 ± 0.03 (n = 100) for WT) (Fig. 5G) and increased Gini coefficient (0.09 ± 0.02 (n = 115) versus 0.08 ± 0.02 (n = 100) for WT) (Fig. 5H). We conclude that deletion of *pcdA* results in the relatively uneven distribution of YSIRK-containing surface adhesins, which results in the altered interaction with host fibrinogen and ultimately impaired SAC formation due to inability to construct the fibrin pseudocapsule.

**Figure 5.**
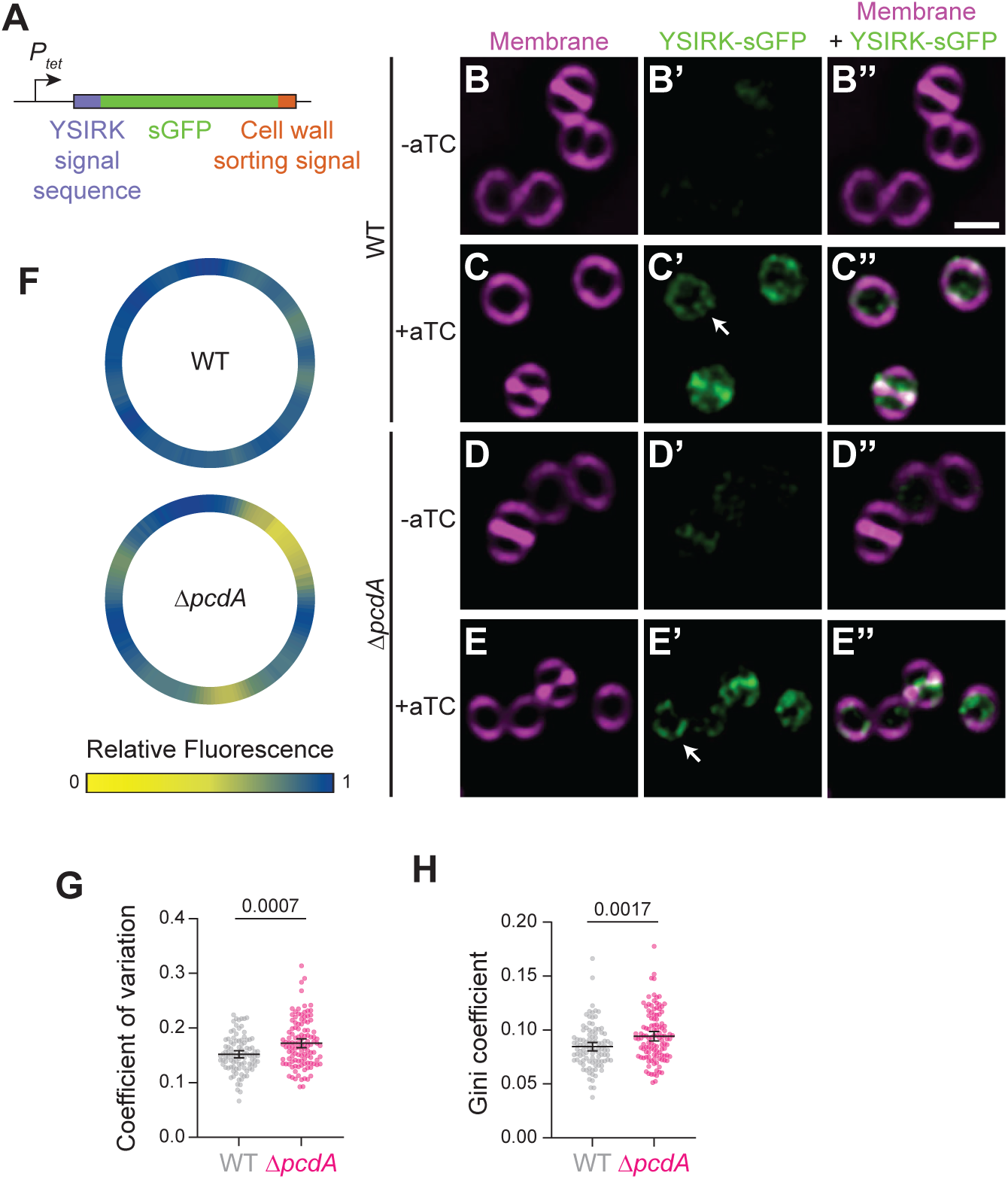
Deletion of *pcdA* impairs the cell surface distribution of YSIRK signal sequence-containing proteins. (A) Schematic of the reporter used to analyze the cell surface distribution of YSIRK signal sequence-containing proteins. The reporter is expressed from an anhydrotetracycline (aTC)-inducible promoter (*P_tet_*) and consists of a YSIRK-containing N-terminal signal sequence (purple), fused to superfolder GFP (sfGFP) (green), and a C-terminal cell wall anchoring signal (sorting signal, orange). (B-E’’) Micrographs of (B-C’’) WT or (D-E’’) Δ*pcdA* cells grown in the absence (B-B’’, D-D’’) or presence (C-C’’, E-E’’) of aTC. (B-E) Cell visualized using membrane stain FM4-64 (magenta); (B’-E’); fluorescence from YSIRK-sGFP (green); (B’’-E’’) overlay, membrane and sGFP. Arrows indicate individual cells whose peripheral fluorescence was quantified in (F) Scale bar, 1 µm. (F) Donut heat map of YSIRK-GFP fluorescence along the perimeter of an individual representative WT and Δ*pcdA* cell. Intensity values were mapped to a continuous yellow-to-blue color scale (yellow = 0; blue = 1). (G-H) Quantification of (G) the coefficient of variation or (H) Gini coefficient of YSIRK-sGFP fluorescence intensity around the circumference of individual WT (gray, n = 100) and Δ*pcdA* (pink, *n* = 115) cells. Bars represent mean ± 95% confidence interval. *P* values were calculated using a Mann-Whitney test.

## DISCUSSION

In this study, we identify orthogonal cell division as an important determinant of staphylococcal abscess community (SAC) formation. Although *S. aureus* was known to divide using consecutive orthogonal planes ^4,32^, the functional consequences of this behavior have remained largely unexplored. We report that disrupting division plane selection through loss of the positive regulator of cell division PcdA results in defects during infection *in vivo* and impaired SAC formation in a three-dimensional *in vitro* model. Together, these findings establish a direct link between the bacterial cell cycle and virulence. We propose a model in which orthogonal cell division promotes an even distribution of YSIRK signal peptide-containing surface adhesins, thereby enabling productive interactions with extracellular matrix components and supporting assembly of the fibrin pseudocapsule that protects SACs in host tissues (Figure 6).

**Figure 6.**
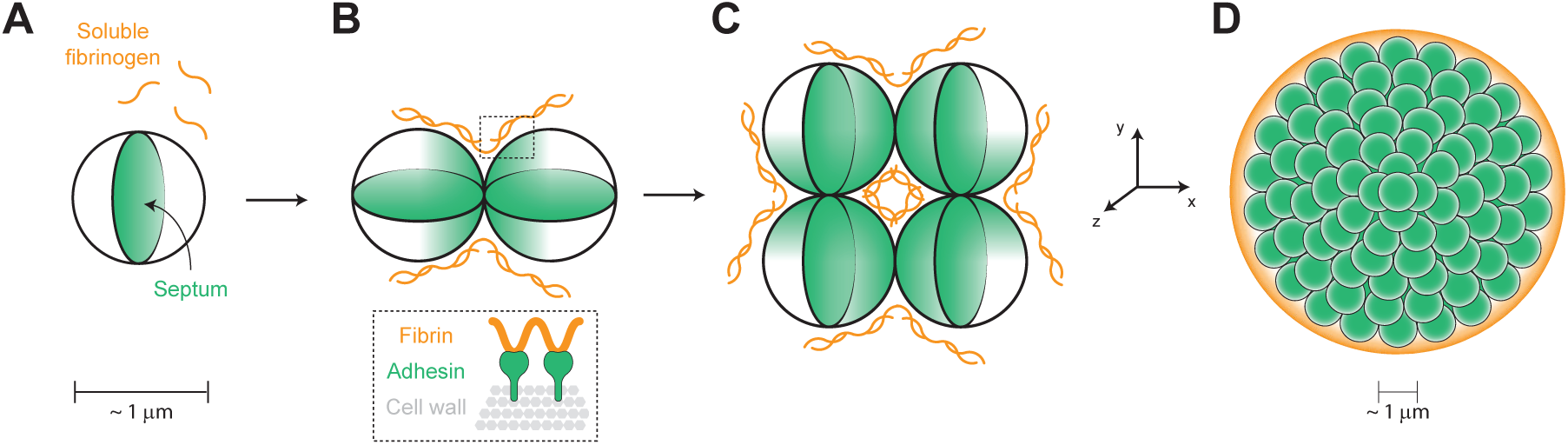
Model for the orthogonal cell division-driven distribution of cell wall anchored surface proteins in a SAC. (A) Spherical staphylococcal cells divide by binary fission, at which time cell wall-anchored surface proteins are incorporated into the newly synthesized septal cell wall (green). Concurrently, bacterial production of coagulases promotes the conversion of soluble fibrinogen into fibrin fibers (orange). (B) YSIRK-containing adhesins involved in fibrinogen and fibrin binding (such as ClfA, FnbpA, or FnbpB), which are enriched at the new cell wall of daughter cells, mediate interactions with the surrounding fibrin network (highlighted box). (C) Successive division in orthogonal planes results in an even distribution of these adhesins over the bacterial cell surface and promotes uniform binding of fibrin. (D) This division pattern promotes the coordinated assembly of a fibrin pseudocapsule surrounding the community, as well as the formation of an interconnected fibrin network within the SAC, thereby supporting three-dimensional SAC growth.

Unlike rod-shaped bacteria, *S. aureus* grows by inserting newly synthesized cell wall at discrete perpendicular planes, imposing a unique spatial organization on the cell surface ^5,33^. Our data indicate that this division geometry plays a central role in patterning surface-exposed virulence factors, particularly adhesins carrying YSIRK signal sequences that are preferentially secreted at the division septum ^20^. We propose that successive orthogonal division events act to redistribute septum-localized proteins over the entire cell envelope, preventing local enrichment or depletion. This process is lost in Δ*pcdA* cells, leading to heterogenous presentation of fibrin-binding adhesins and impaired interactions with the extracellular fibrin matrix. These observations suggest that orthogonal cell division coordinates cell cycle progression with cell surface organization, a feature that is critical for *S. aureus* adaptation and persistence within host environments.

Other factors contribute to division plane selection in *S. aureus*, including the generation of transient polarity due to subtle elongation mediated by PBP3-RodA ^7^ and the nucleoid occlusion protein Noc ^9^. Interestingly, though, only mutations affecting the elongation machinery impair SAC development, suggesting that cell wall-dependent pathways are particularly relevant for this mode of growth, whereas chromosome-dependent pathways may play a lesser role, at least under the conditions tested here. Nevertheless, Noc has been shown to influence initiation of DNA replication ^8^, and how coordination of chromosome dynamics with cell division contributes to SAC formation remains unstudied. Further work will be needed to determine if these pathways intersect during growth in a host.

Disruption of orthogonal division does not increase uptake of individual bacteria by phagocytic cells. This observation is consistent with previous studies showing that *S. aureus* clumping, rather than single-cell properties, plays a key role in evading phagocytosis during early stages of infection ^34^. The contribution of fibrinogen-mediated cellular aggregation to immune evasion is well established ^35,36^, and our findings suggest that defects in SAC architecture, rather than altered interactions at the level of individual cells, likely underlie the in vivo phenotype observed for the Δ*pcdA* strain. Thus, while surface defects caused by loss of PcdA may be largely tolerated in isolated bacteria *in vitro*, they have consequences for collective behavior and virulence in host environments.

Recent work highlights the contribution of the cell wall and cell division to *S. aureus* pathogenesis, including regulation of virulence factor sorting ^37^ and the requirement for controlled cell wall hydrolysis during cell separation to ensure proper display of surface-anchored proteins ^38,39^. Loss of PcdA alters the spatial patterning of fibrinogen binding and YSIRK-containing proteins across the cell surface without abolishing their overall exposure, suggesting that organization, rather than simply abundance, is the primary determinant of the observed phenotypes. Importantly, infection-relevant conditions have also been shown to induce remodeling of the cell wall. For example, *S. aureus* isolated from infected tissues displayed increased cell wall thickness and reduced crosslinking ^40^, and exposure to serum similarly promoted cell wall thickening ^41,42^. These observations are therefore consistent with the idea that dynamic changes in cell wall architecture during infection may influence surface protein organization and, consequently, bacterial interactions with host factors.

Taken together, our findings highlight the importance of studying bacterial cell division in host-relevant contexts, where the consequences of cell cycle regulation extend beyond growth and morphology to directly shape collective behaviors and virulence. By integrating *in vivo* infection models with *in vitro* conditions that recapitulate key features of the host, our work reveals how division geometry governs surface organization and productive interactions with fibrin that are essential for SAC development. More broadly, these results emphasize the extracellular matrix as a critical interface in *S. aureus* pathogenesis and support the idea that disrupting bacterial-matrix interactions, or the cellular pathways that control them, represents a promising strategy to weaken bacterial communities and enhance host clearance ^30,43,44^.

## ACKNOWLEDGEMENTS

We thank Susan Gottesman and Anupama Khare for discussions, members of the K.S.R. lab for comments on the manuscript, and Yuen Chang for technical advice and the RAW264.7 cell line. This research was partially supported by funding from the Burroughs Wellcome Fund (1180500) to K.M.D. and funding from the National Institutes of Health (NIH; T32AI007417) to P.G, and the Intramural Research Program of the NIH, National Cancer Institute, Center for Cancer Research (K.S.R.). The contributions of the NIH authors are considered Works of the United States Government. The findings and conclusions presented in this paper are those of the authors and do not necessarily reflect the views of the NIH or the U.S. Department of Health and Human Services.

## DECLARATION OF INTERESETS

The authors declare no competing interests.

## EXPERIMENTAL PROCEDURES

### Ethics statement and murine infection model

All animal experiments were carried out as previously described ^14^. Protocols were approved by the Johns Hopkins University Institutional Animal Care and Use Committee (protocol MO23H310). *S. aureus* strains were prepared in sterile PBS to 10^6^ CFU in 100 µl for inocula. 6-to 8-week-old female C57BL/6 mice (Jackson Laboratories) under isoflurane anesthesia were inoculated intravenously via the rertro-orbital route. At 3 days post-inoculation, mice were euthanized and kidneys were harvested and fixed in 4% PFA overnight at 4 ⁰C for histology.

### Strains and culture conditions

Strains and plasmids used in this study are listed in Table S1-S2. *Escherichia coli* strains were grown in Lysogenic Broth (LB) (KD Medical) and LB agar. The medium was supplemented with 100 μg ml^-1^ spectinomycin for selection and plasmid maintenance as required. *Staphylococcus aureus* strains used in this study are derivatives of JE2, a plasmid-less derivative of the methicillin-resistant *S. aureus* USA300 lineage ^45^. *S. aureus* was grown in Tryptic Soy Broth (TSB), Modified M63 medium ^46^ (13.6 g L^-1^ KH_2_PO_4_, 2 g L^-1^ (NH_4_)_2_SO_4_, 0.8 µM ferric citrate, 1 mM MgSO_4_; pH adjusted to 7 using KOH) supplemented with 0.3% glucose, 1x ACGU solution (Teknova), 1x supplement EZ (Teknova), 0.1 ng L^-1^ biotin and 2 ng L^-1^ nicotinamide, or Roswell Park Memorial Institute (RPMI) 1640 medium without phenol red and supplemented with 2 g L^-1^ sodium bicarbonate (Sigma). When required, medium was supplemented with 5 μg ml^-1^ erythromycin, 10 μg ml^-1^ chloramphenicol or 1.5 μg ml^-1^ tetracycline. For growth on solid media, Tryptic Soy Agar (TSA) was used, supplemented with antibiotics at the indicated concentrations as needed.

### Construction of reporter strains

To construct strains expressing mScarlet3 from the *rplM* promoter, DNA encoding codon-optimized mScarlet3 was amplified from a synthetic gBlock (Integrated DNA Technologies) using primers 432 and 433. The upstream region of *rplM* was amplified from *S. aureus* genomic DNA using primers 430 and 436. The resulting fragments were assembled into pPhi11 ^5^, digested with *BamH*I and *EcoR*I using Gibson assembly (New England Biolads), yielding plasmid pFRL241.

To construct strains expressing a YSIRK-containing sGFP reporter under the control of an anhydrotetracycline-inducible promoter, the *tet* promoter was amplified from pJB38 ^47^ using primers 465 and 466. The DNA fragment encoding the YSIRK signal sequence was amplified from *S. aureus* genomic DNA using primers 467 and 468. The *sGFP* gene was amplified using primers 469 and 470 from pFRL126 ^5^, and a fragment encoding the cell wall sorting signal was amplified from gDNA using primers 471 and 472. All fragments were assembled into pPhi11 digested with *BamH*I and *EcoR*I using Gibson assembly, generating plasmid pFRL251.

Plasmids were initially introduced into the restriction-deficient *S. aureus* strain RN4220 ^48^ carrying plasmid pLL2757, which encodes the φ11 integrase ^49^. Integration at the φ11 *attB* site was selected on TSA plates supplemented with 1.5 μg ml^-1^ tetracycline. Integrated constructs were subsequently transduced into the desired strain backgrounds using bacteriophage φ85.

### Fluorescence microscopy of mouse tissues

Preparation of kidneys for imaging was done as described previously ^14^. Briefly, kidneys were fixed in 4% PFA overnight, frozen-embedded in O.C.T. compound (Tissue-Tek, BWR), and then cut in 10 μm sections using a cryostat microtome (Microm HM 505e). Sections were mounted on charged microscope slides (HistoBond, VWR) and then thawed in PBS at room temperature. Nuclei were stained by incubation Hoechst (1:10,000 dilution in PBS) for 15 min. Samples were imaged using a Zeiss Axio Observer 7 inverted fluorescent microscope with a 63x oil objective and Apotome 2 equipped with a Axiocam 702 mono camera (Zeiss). Images were processed using ZEN3.10 software.

### Uptake by RAW264.7 murine macrophages

RAW264.7 cells were maintained in DMEM supplemented with 10% fetal bovine serum (FBS) at 37 ⁰C in a humidified atmosphere containing 5% CO₂. The day before infection, RAW264.7 cells were gently detached from culture dishes using a cell scraper, collected by centrifugation, and resuspended in fresh pre-warmed DMEM with 10% FBS. Cells were counted, and 2.5 × 10^6^ cells were seeded onto 35-mm dishes containing glass coverslips. Dish was filled with 2 ml of DMEM with 10% FBS and incubated overnight. After 24 h, the dishes contained approximately 5 × 10^6^ cells, resulting in near-confluent monolayers.

*S. aureus* strains expressing mScarlet3 under the control of the *rplM* promoter were grown overnight in modified M63 medium. Overnight cultures were diluted to an initial OD_600_ of 0.1 in fresh modified M63 medium and incubated at 37 ⁰C with shaking until reaching an OD_600_ of 0.5, corresponding to ∼3 × 10^8^ CFU ml^-1^. A volume containing 2.5 × 10^7^ CFU was harvested, washed three times with DMEM, and resuspended in 2 ml of DMEM supplemented with 2% FBS. The medium from the macrophage dishes was aspirated and replaced with the bacterial suspension. Plates were incubated for 2 h at 37 ⁰C and 5% CO₂ to allow phagocytosis. Following infection, the supernatant was removed, and the cells were washed three times with sterile PBS. To kill extracellular bacteria, cells were incubated for 1 h at 37 ⁰C in DMEM containing 10% FBS, 100 µg ml gentamicin^-1^, and 50 µg ml^-1^ lysostaphin.

For microscopy, after antibiotic treatment, the medium was aspirated, and cells were washed twice with 2 ml of sterile PBS. Cells were then incubated for 15 min at room temperature in the dark with 1 ml of pre-warmed staining solution (RPMI without phenol red containing 1 µg ml^-1^ Hoechst 33342 and 2 µg ml^-1^ WGA-488). The staining solution was removed, and cells were washed twice with pre-warmed RPMI. Finally, 1 ml of RPMI without red phenol red was added, and cells were imaged using a DeltaVision inverted microscope as described previously ^50^.

For flow cytometry, the same infection and antibiotic treatment protocol was followed, except cells were seeded in 35-mm dishes without coverslips. After antibiotic treatment, cells were washed twice with 2 ml of sterile PBS, resuspended in 1 ml of PBS, and gently detached using a cell scraper. The suspension was transferred to sterile microtubes. Cells were stained with Zombie Aqua Fixable Viability Kit (BioLegend) according to the manufacturer’s instructions to discriminate viable from non-viable cells. After 20 min incubation at room temperature in the dark, cells were collected by centrifugation at 200 × g for 5 min at 4 ⁰C, resuspended in 1 ml PBS, and analyzed by flow cytometry using BD FACSymphony A5 (BD Biosciences). All data were analyzed using FlowJo 11.1 (Becton Dickinson and Company).

### Formation of staphylococcal abscess-like communities in 3D collagen gels

Growth in collagen gels was adapted from a previously reported protocol ^16^ with several modifications. *Staphylococcus aureus* strains were cultured overnight at 37 ⁰C with shaking in RPMI medium lacking phenol red. The following day, cultures were diluted into fresh RPMI without phenol red and incubated at 37 ⁰C with shaking until reaching an OD_600_ of 0.5, corresponding to approximately 3 × 10^8^ CFU ml^-1^. Bacterial cultures were then diluted to an OD_600_ of 0.01 in RPMI medium without phenol red supplemented with 1.7 mg ml^-1^ rat tail collagen I (Cellink). A volume of 100 µl was added to each well of a glass-bottom μ-slide 8-well chamber slide (Ibidi), ensuring even distribution across the well surface. Slides were incubated for 45 minutes at 37 ⁰C in a humidified incubator with 5% CO_2_ to allow gelation. After that, 300 μL of FBN/PT medium (RPMI supplemented with 3 mg ml^-1^ human fibrinogen, 30 µg ml^-1^ prothrombin, and 10% fetal bovine serum) was gently added to each well. Slides were incubated overnight at 37 ⁰C with 5% CO_2_. Staphylococcal abscess-like communities were imaged after 16-18 h of incubation using a DeltaVision fluorescence microscope.

### Generation of stable EGFP-expressing RAW264.7 macrophages

RAW264.7 murine macrophages cells were stably transfected to express enhanced green fluorescent protein (EGFP) using the PiggyBac transposon system. The donor plasmid pBP-EF1α-EGFP-T2A-Puro (Addgene plasmid #133400), encoding EGFP and a puromycin resistance cassette, was co-transfected with the helper plasmid PB210PA-1 (System Biosciences), which expresses the PiggyBac transposase. Cells were maintained in Dulbecco’s Modified Eagle Medium (DMEM) supplemented with 10% fetal bovine serum and 1% penicillin-streptomycin at 37 ⁰C with 5% CO_2_. For transfection, cells were seeded in 100-mm tissue culture dishes and grown to ∼70–80% confluency. A total of 10 μg of donor plasmid and 5 μg of transposase plasmid were diluted in JetPrime® buffer (Polyplus Transfection, Illkirch, France), mixed with JetPrime® reagent, and incubated at room temperature for 10 minutes before being added dropwise to the cells. 4 h after transfection, the medium was replaced with fresh complete DMEM. At 48 h post-transfection, cells were subjected to selection with 8 μg ml^-1^ puromycin. Selection medium was refreshed every 2-3 days for 7-10 days until non-transfected control cells were eliminated and a stable EGFP-expressing population was established. EGFP expression was confirmed by fluorescent microscopy.

### Coagulase assay

*S. aureus* strains were grown overnight in modified M63 medium. Overnight cultures were used to inoculate 1 ml of modified M63 medium supplemented with 3 mg ml^-1^ fibrinogen and 30 µg ml^-1^ prothrombin to an initial OD_600_ of 0.01. Cultures were incubated statically at 37 °C for 16 h to allow coagulase-mediated conversion of fibrinogen to fibrin. Coagulation was assessed visually by the formation of a solid gel that remained intact upon gentle inversion of the tube.

### Soluble fibrinogen binding assays

*S. aureus* strains were grown overnight in modified M63 medium. Cultures were diluted the following day to an initial OD_600_ of 0.1 and grown at 37°C with shaking until reaching an OD_600_ of 0.5. Cells from 500 μl were then harvested by centrifugation and resuspended in 100 μl of phosphate-buffered saline (PBS) containing 0.25 mg ml^-1^ fibrinogen conjugated to Alexa Fluor 647 (Thermo Fisher Scientific). Samples were incubated at room temperature for 5 min, after which cells were collected by centrifugation and washed three times with PBS. Cells were then imaged using a DeltaVision fluorescence microscope.

### Analysis of YSIRK-containing reporter

*S. aureus* strains carrying plasmid pFRL251 were grown in modified M63 medium. Overnight cultures were diluted into fresh modified M63 to an initial OD_600_ of 0.1 and grown at 37 °C with shaking until reaching an OD_600_ of 0.4. Reporter expression was induced by addition of anhydrotetracycline to a final concentration of 300 ng ml^-1^, and cultures were incubated for an additional 90 min. Cells were then harvested by centrifugation and resuspended in PBS containing 50 μg ml^-1^ FM4-64 for membrane staining. Cells were imaged using a DeltaVision fluorescence microscope. Image analysis was performed using ImageJ version 1.54p. To generate donuts heatmaps, line profiles were manually traced along the cell boundary using ImageJ, and fluorescence intensity values were extracted along the traced path. The resulting intensity was normalized to the maximal signal within each cell. Normalized intensity values (0-1) were converted into a continuous color scale ranging from yellow (low intensity) to blue (high intensity). Color values were assigned by linear interpolation between the two endpoints in RGB space.

## SUPPLEMENTAL INFORMATION

**Figure S1.**
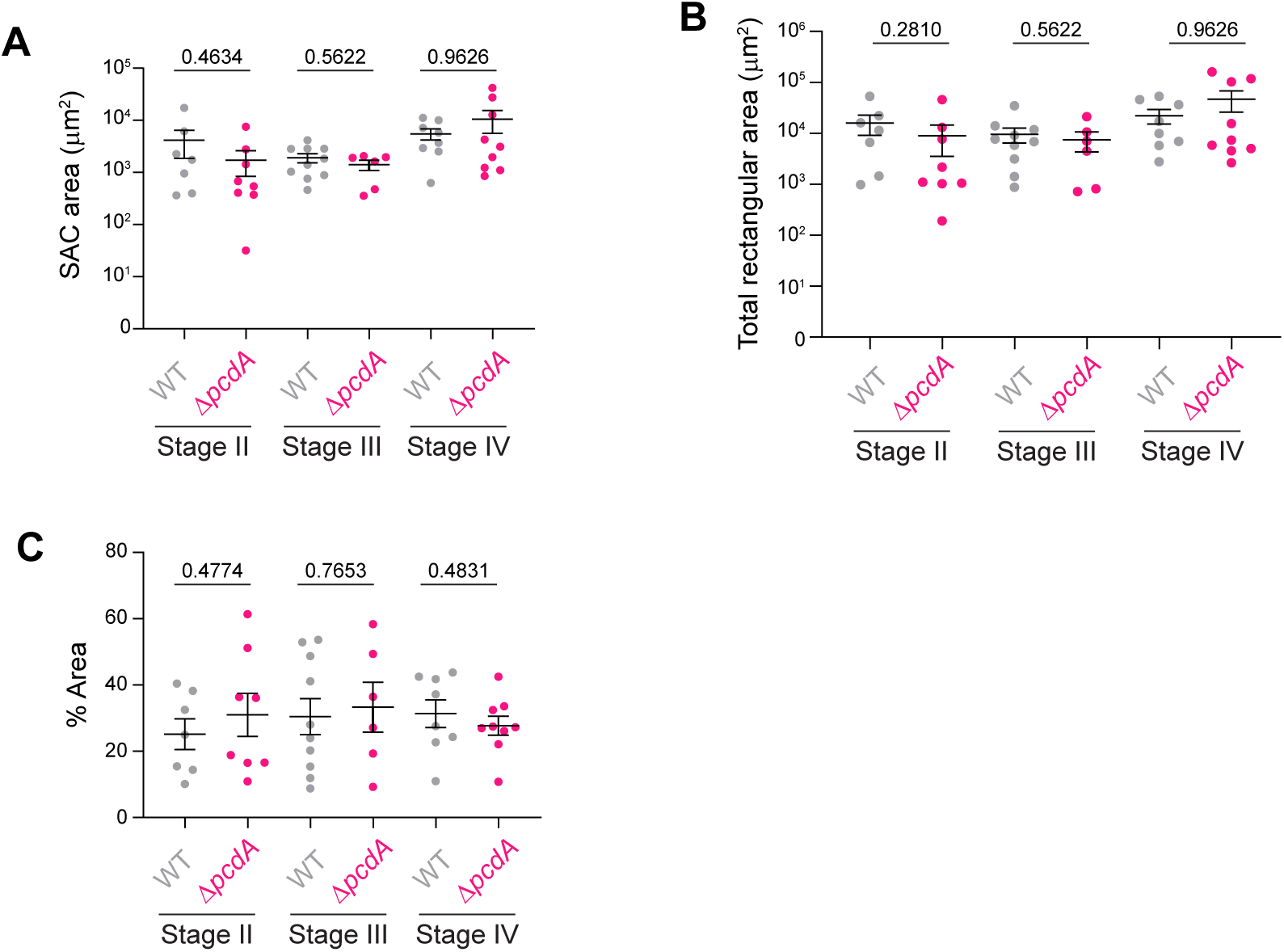
Characterization of different stages of staphylococcal abscess communities. (A) Area (μm^2^) containing bacteria in stage II, stage III, and stage IV lesions for WT (grey) and Δ*pcdA* (pink). (B) Quantification of the total rectangular area (μm^2^) of stage II, stage III, and stage IV lesions in kidneys infected with WT (grey) and Δ*pcdA* (pink). The total rectangular area represents the area of a best-fit rectangle around the bacterial cells. (C) Percentage of the stage II, stage III, and stage IV lesion area occupied by bacteria in kidneys infected by WT (grey) and Δ*pcdA* (pink). For all graphs, bars indicate the mean ± standard error of the mean. *P* values were determined using a Mann-Whitney test.

**Figure S2.**
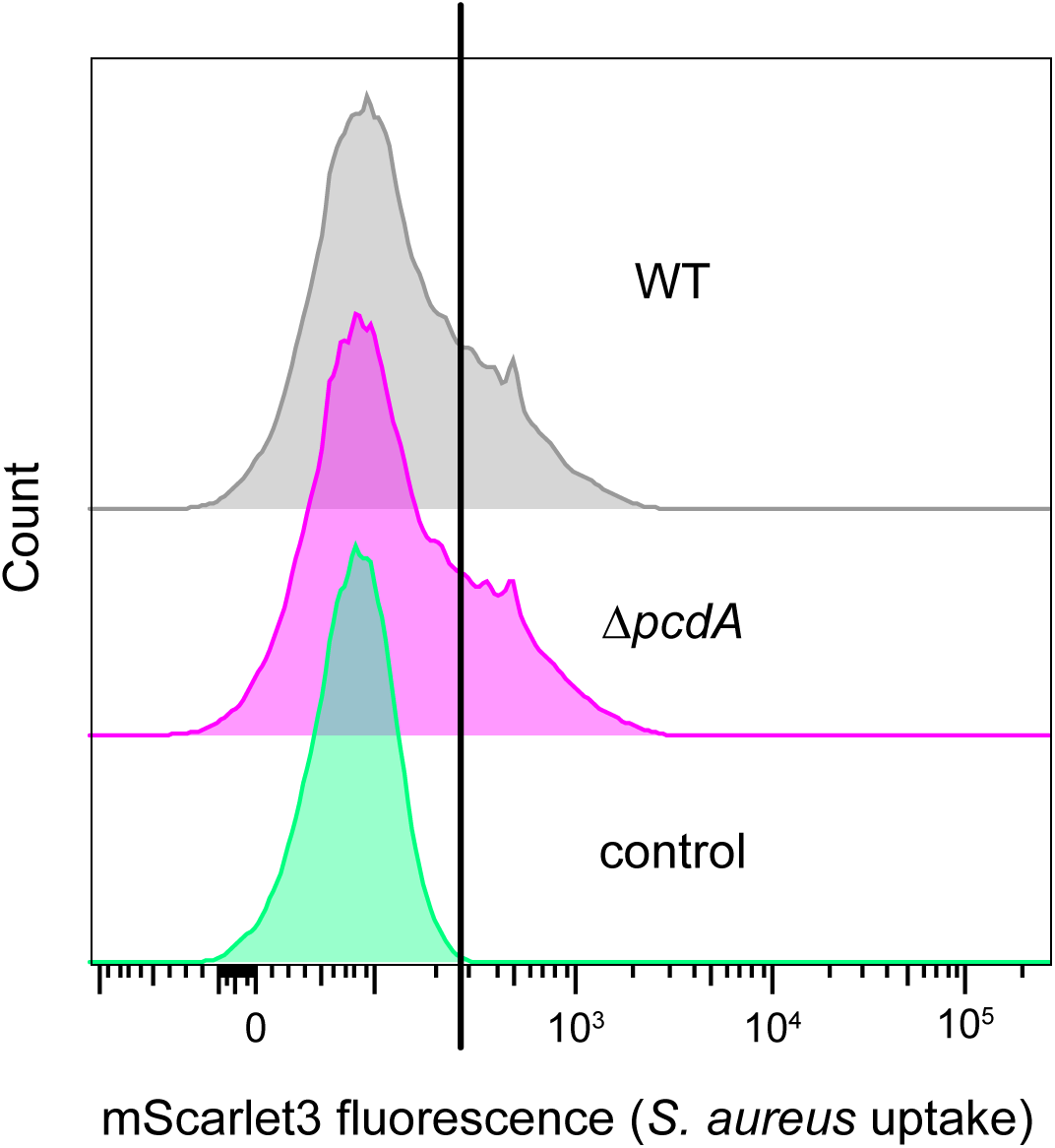
Representative flow cytometry histograms of RAW264.7 macrophages incubated with *S. aureus*. Histograms show the distribution of mScarlet3 intensity in RAW264.7 cells following incubation with WT (grey; n = 46,073) or Δ*pcdA* (pink; 45,390) strains, or in control samples lacking bacteria (green; n = 31,775), which were used for normalization. The *y*-axis represents cell counts, and the *x*-axis represents mScarlet3 fluorescence intensity.

**Figure S3.**
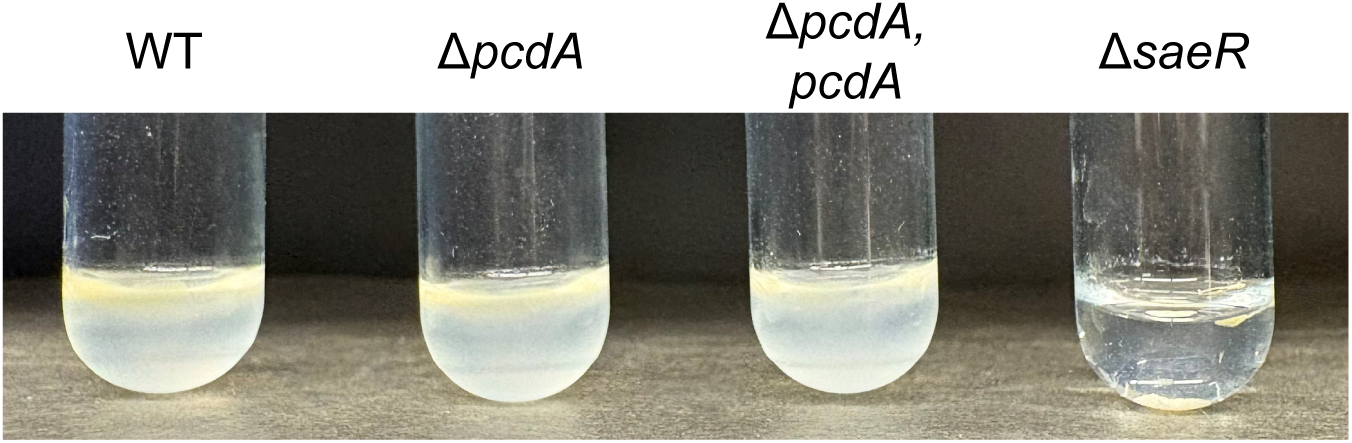
Deletion of *pcdA* does not affect fibrinogen-to-fibrin coagulation. Representative photograph of test tubes containing the indicated *S. aureus* strains grown overnight in the presence of fibrinogen and prothrombin to allow coagulation. Samples were imaged the following day. The Δ*saeR* strain was included as a negative control due to its inability to promote fibrinogen coagulation.

**Table S1.**
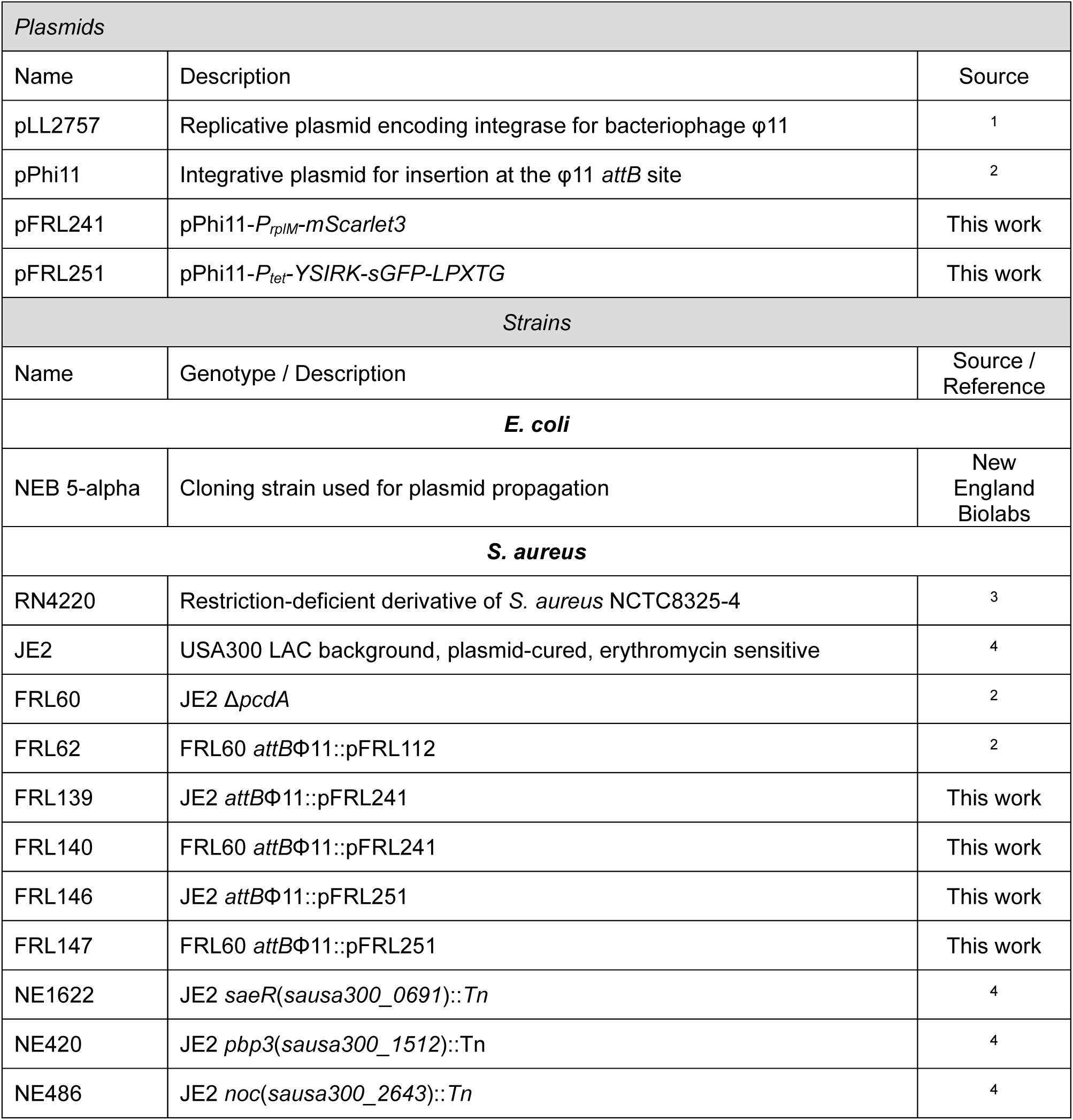
Plasmids and strains used in this study.

| *Plasmids* |  |  |
| --- | --- | --- |
| Name | Description | Source |
| pLL2757 | Replicative plasmid encoding integrase for bacteriophage ϕ11 | 1 |
| pPhi11 | Integrative plasmid for insertion at the ϕ11 *attB* site | 2 |
| pFRL241 | pPhi11-*P<sub>rpIM</sub>-mScarlet3* | This work |
| pFRL251 | pPhi11-*P<sub>tet</sub>-YSIRK-sGFP-LPXTG* | This work |
| *Strains* |  |  |
| Name | Genotype / Description | Source / Reference |
| *E. coli* |  |  |
| NEB 5-alpha | Cloning strain used for plasmid propagation | New England Biolabs |
| *S. aureus* |  |  |
| RN4220 | Restriction-deficient derivative of *S. aureus* NCTC8325-4 | 3 |
| JE2 | USA300 LAC background, plasmid-cured, erythromycin sensitive | 4 |
| FRL60 | JE2 Δ*pcdA* | 2 |
| FRL62 | FRL60 *attB*Φ11::pFRL112 | 2 |
| FRL139 | JE2 *attB*Φ11::pFRL241 | This work |
| FRL140 | FRL60 *attB*Φ11::pFRL241 | This work |
| FRL146 | JE2 *attB*Φ11::pFRL251 | This work |
| FRL147 | FRL60 *attB*Φ11::pFRL251 | This work |
| NE1622 | JE2 *saeR*(*sausa300_0691*)::Tn | 4 |
| NE420 | JE2 *pbp3*(*sausa300_1512*)::Tn | 4 |
| NE486 | JE2 *noc*(*sausa300_2643*)::Tn | 4 |

**Table S2.**
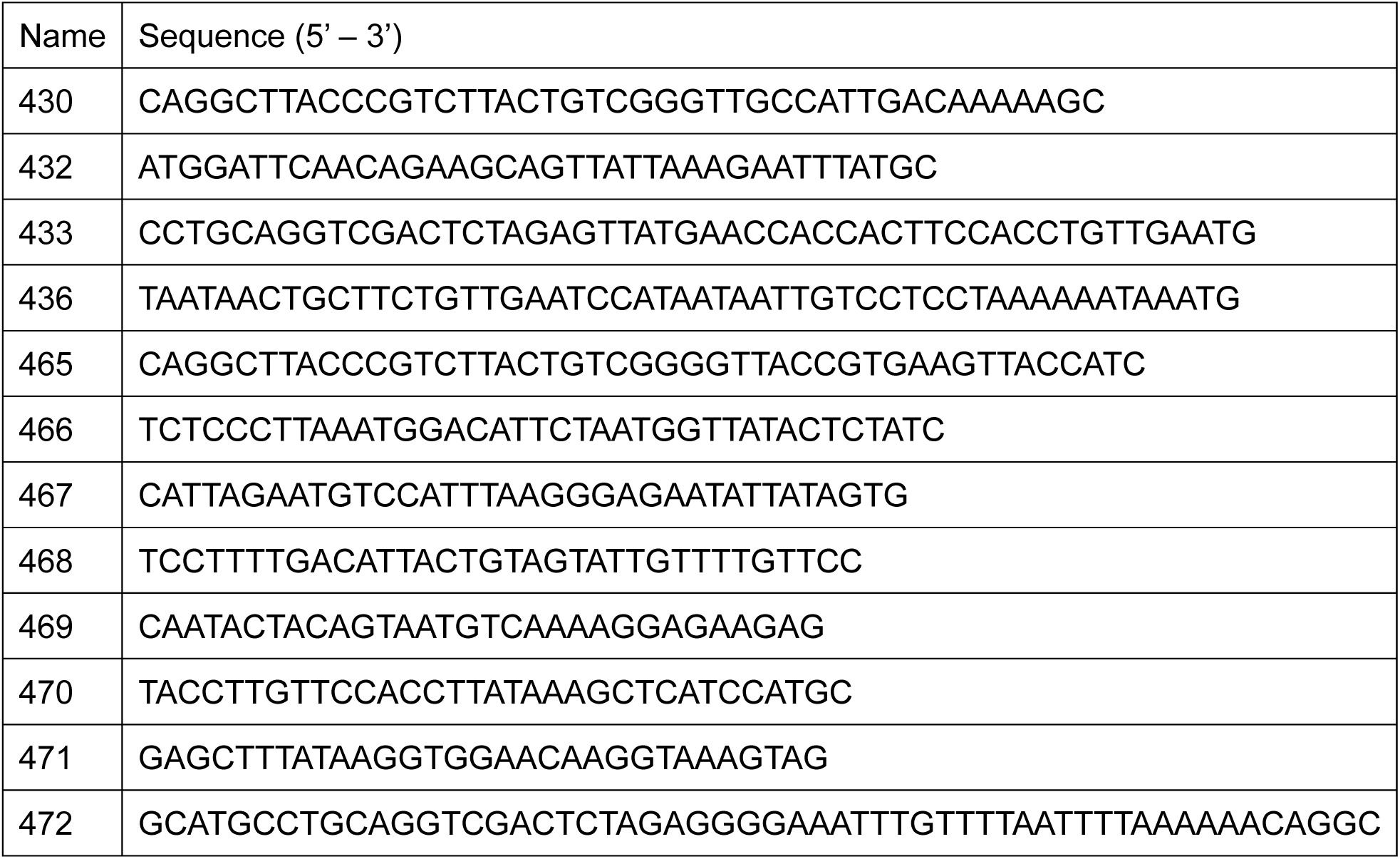
Oligonucleotides used in this work.

